# Bursting on a two state stochastic model for gene transcription in *Drosophila* embryos

**DOI:** 10.1101/107979

**Authors:** Romain Yvinec, Luiz Guilherme S. da Silva, Guilherme N. Prata, John Reinitz, Alexandre Ferreira Ramos

**Affiliations:** Physiologie de la Reproduction et des Comportements, Institut National de la Recherche Agronomique (INRA) UMR85, CNRS-Université François-Rabelais UMR7247, IFCE, Nouzilly, 37380 France; Escola de Artes, Ciências e Humanidades & Núcleo de Estudos Interdisciplinares em Sistemas Complexos, Universidade de São Paulo, Av. Arlindo Béttio, 1000 CEP 03828-000, São Paulo, SP, Brazil; Department of Statistics, Department of Ecology and Evolution, Department of Molecular Genetics and Cell Biology, University of Chicago 5734 S. University Avenue Eckhart 134 Chicago, IL 60637, USA and; Departamento de Radiologia – Faculdade de Medicina, Universidade de São Paulo & Instituto do Câncer do Estado de São Paulo, São Paulo, SP CEP 05403-911

## Abstract

Recent experimental data on the transcription dynamics of *eve* gene stripe two formation of *Drosophila melanogaster* embryos occurs in bursts of multiple sizes and durations. That has motivated the proposition of a transcription model having multiple ON states for the promoter of the *eve* gene each of them characterized by different synthesis rate. To understand the role of multiple ON states on gene transcription we approach the exact solutions for a two state stochastic model for gene transcription in *D. melanogaster* embryos and derive its bursting limit. Simulations based on the Gillespie algorithm at the bursting limit show the occurrence of bursts of multiple sizes and durations. Based on our theoretical approach, we interpret the aforementioned experimental data as a demonstration of the intrinsic stochasticity of the transcriptional processes in fruit fly embryos. Then, we conceive the experimental arrangement to determine when gene transcription has multiple ON promoter state in a noisy environment.

---

The control of transcription in cells is characterized by intrinsic fluctuations which result from the small number of molecules of proteins which control this process. In prokaryotes, the stochastic theory for this process is relatively well developed because of good understanding of the fundamental reaction mechanisms, but in eukaryotes the detailed reaction and control mechanisms are poorly understood, and a detailed stochastic theory does not exist. Nevertheless, high resolution experimental studies clearly reveal intrinsic fluctuations in the form of bursts of transcription of variable size and duration [1–8]. The number of underlying transcriptional states implied by this bursting behavior remain unclear. Previous investigations have explained the variable size and duration of bursts in terms of the random initiation of multiple underlying states [9], while variation in the duration of bursts has been explained by a two state stochastic model [10]. In the latter study an *ansatz* required by the experimental data prevented estimation of burst size, while the former used a deterministic model of the binding and elongation of RNA PolII. In this manuscript we consider the exact solutions for the two state stochastic model to calculate its burst limit approximation rigorously and show the occurrence of variable burst size using Gillespie’s exact simulation algorithm. This provides an estimate of the average burst size in one previous study [10] and demonstrates that the data reported in the other study [9] is fully compatible with a two state model.

Here we use a two state master equation model of transcription which has exact solutions given by confluent hypergeometric functions [11,12] which has been previously applied to the control of the transcription of *even-skipped* stripe 2 in *D. melanogaster* [13]. The stochastic variables of the model are the number of mRNA molecules, denoted by *n*, and the state of the gene’s promoter being ON or OFF. The probability for the promoter state to be ON (or OFF) at time *t* when *n* mRNA molecules are found within a nucleus is denoted by *α*_*n*_(*t*) (or *β*_*n*_(*t*)). mRNA synthesis occurs at rate *k* when the gene is ON and is zero when the gene is OFF. mRNA degradation occurs at rate *ρ*. The promoter transition from the ON to the OFF (or from the OFF to the ON) state has rate *h* (or *f*). We omit the temporal dependence of the probabilities such that *α*_*n*_(*t*) ≡ *α*_*n*_ and *β*_*n*_(*t*) ≡ *β*_*n*_ and write the maste equation for the steady state limit as

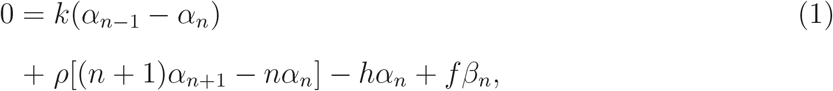

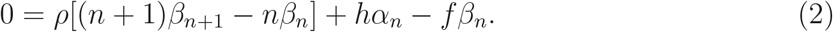

The steady state probabilities of finding *n* mRNA molecules independently of the gene state are denoted by *ϕ*_*n*_, with *ϕ*_*n*_ = *α*_*n*_ + *β*_*n*_, given by

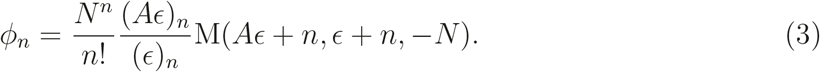

M(*a, b, z*) denotes the KummerM function and (*x*)_*n*_ = *x*(*x* + 1)… (*x* + *n* − 1) denotes the Eq. (3) is written in terms of the parameters (*N, ε, A*) defined by

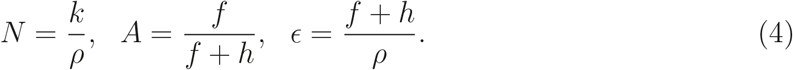

*N* denotes the expectation of the number of mRNA molecules when the promoter is exclusively ON. The steady state probability of finding the promoter in the ON state is denoted by *A*, with 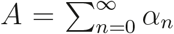. ε gives the ratio of the switching rate between ON and OFF to the rate of mRNA degradation. Thus, *ε*≫ 1 implies that the gene switches multiple times during the mean lifetime of an mRNA molecule, while *ε* ≪ 1 implies that the gene stays ON or OFF for a time that is longer than the lifetime of the message. Of the two experimental studies mentioned above, one [10] reports data for which *A* ∼ 0, while the other reports data where *ε* ≫ 1. We consider each case in turn below.

*I. Bursting limit.* In the work of Suter *et al.*[10], Fig. 2A shows that the number of mRNA molecules varies from 0 to 8, and is frequently 0. This indicates that *A* ∼ 0. We now show that in this limit, the steady state solutions for the master equations (1) and (2) in Eq. (3) have the negative binomial probability distribution. Let us evaluate *ϕ*_*n*_ at the limit of *A* ∼ 0 and *ε, N* ≫ 1 with the ratio *δ* = *N/ε* kept as a finite constant. That transforms *ϕ*_*n*_ into 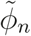 where

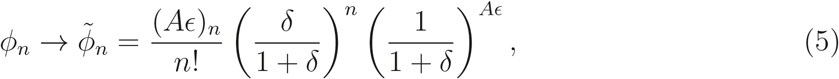
the negative binomial distribution which governs mRNA numbers in transcriptional bursts, as has previously been shown for translational bursts [14,15].

**FIG. 1.**
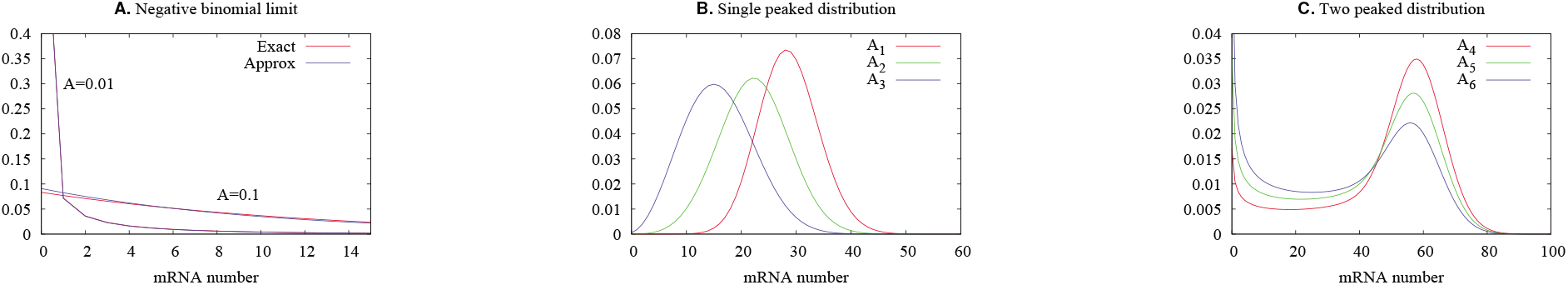
Here we show the probability distributions obtained with Eqs. (3) and (5). The Fano factors for those distributions are given by 1 + *N* (1 − *A*)*/*(1 + *ε*). A. The approximation of Eq. (3) by Eq. (5) with parameters (*N, ε*) = (100, 10) and the value of *A* is given as a key within the graphs. The Fano factors for *A* = 0.01 and *A* = 0.001 are, respectively, 10. and 10.08. Where the red trace is not visble, the curves overlap. B. Single peaked probability distributions of exact solutions of Eq.(3) for the parameters (*N, ε*) = (28.9, 7.2) and (*A*_1_, *A*_2_, *A*_3_) = (0.99, 0.77, 0.55) with the respective Fano factors given by ≈ (1.04, 1.81, 2.59). C. Two-peaked probability distributions of the exact solutions of Eq. (3) for the parameters (N, ε) = (60, 0.75) and (*A*_4_, *A*_5_, *A*_6_) = (0.8, 0.7, 0.6) with the respective Fano factors given by ≈ (7.86, 11.29, 14.71).

*II. Probability distributions.* Fig. (1A) shows a comparison between the probabilities *ϕ*_*n*_ and their approximations 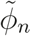 for two sets of parameters (*N, ε, A*), which we denote below as *P*_1_≡ (100, 10, 0.1) and *P*_2_≡ (100, 10, 0.01). The accumulated difference between the two probability distributions, 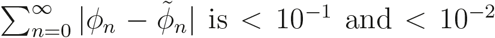, respectively, for the parameter sets *P*_1_ and *P*_2_. Note that the value of *A* indicates the order of magnitude of the accumulated error of the negative binomial approximation. Figs. (1.B) and Fig. (1.C) show single peaked and two-peaked probability distributions of exact solutions of Eq. (3). The two peaked distributions occur when *ε <* 1 with the relative height of the peaks dependent on the value of *A*. The modes of the single peaked distributions approach *N* as *A* approaches 1.

*III. Exact simulations.* Fig. (2) shows three realizations of the stochastic process governed by the probabilities *α*_*n*_(*t*)*, β*_*n*_(*t*). The blue (red) lines show the number of mRNAs (promoter state) as a function of time. Fig. (2.A) shows trajectories for transcriptional bursting in which mRNA numbers at steady state are governed by the negative binomial limit of the probability distribution of the Eq. (3). Fig. (2.B) presents trajectories obtained from transcriptional dynamics characterized by a single peaked probability distribution, as in Fig. (1.B). The bursting of mRNAs occurs during the promoter ON state and the A - Stochastic promoter dynamics and stochastic bursty transcriptional dynamics B - Stochastic promoter dynamics and stochastic bursty transcriptional dynamics C - Stochastic promoter dynamics and stochastic bursty transcriptional dynamics OFF-ON-OFF state switching time duration appears as a single vertical line. The bimodal distribution limit of Fig. (1.C) has its corresponding trajectories shown in Fig. (2.C).

**FIG. 2.**
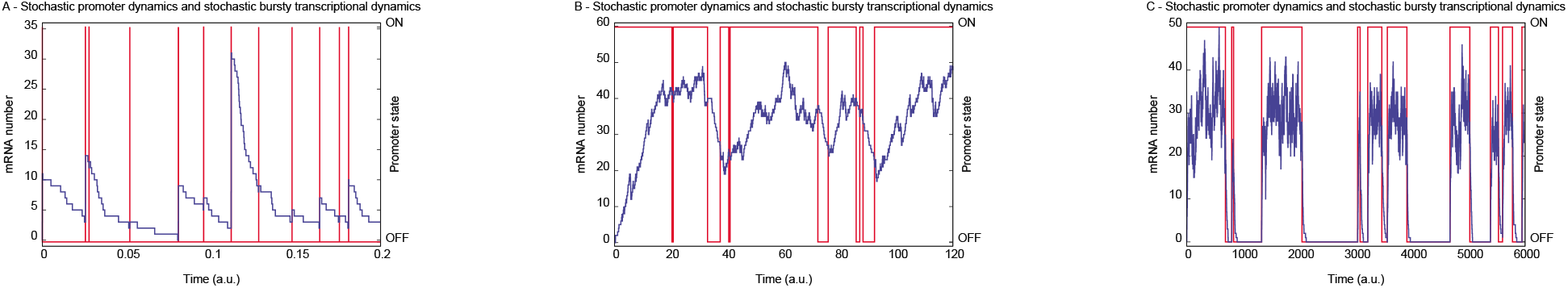
Realizations of the dynamics of the number of mRNAs and corresponding promoter state by the exact simulation algorithm of Gillespie [16] are presented. The left and right side vertical axes show the number of mRNA molecules and promoter state, respectively. The number of mRNA molecules (promoter state) trajectory is indicated in blue (red). The graphs A, B, and C were constructed using parameters (*N, ε, A, ρ*), respectively, equal to (8 × 10^3^, 10^3^, 10^−3^, 1), (40, 5, 0.9, 0.002), and (0.03, 0.1, 0.5, 10^−3^).

**FIG. 3.**
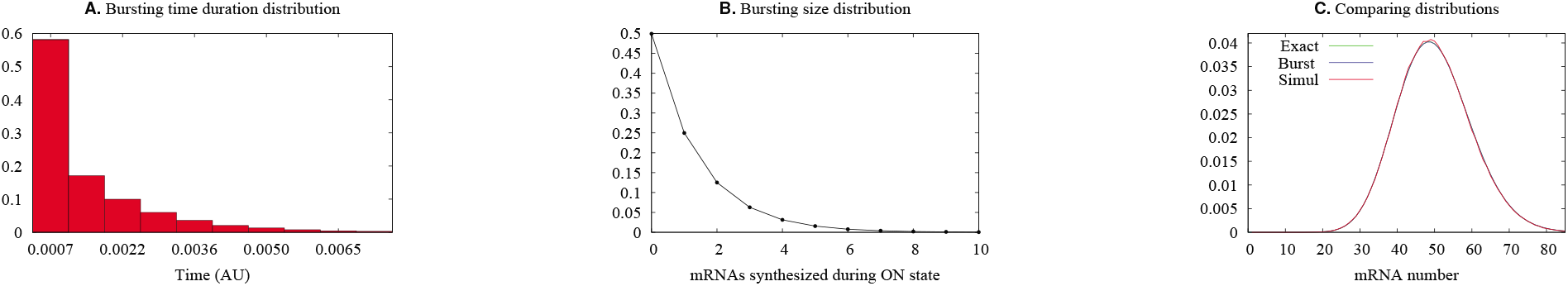
The histograms for the burst duration and burst size for parameters (*N, ε, A*) = (10^4^, 10^4^, 0.005) are presented in A and B, respectively. C shows a comparison of the steady state distribution of the mRNA numbers obtained from Eq. (3), denoted by the label ‘Exact’, the Eq. (5), denoted by ‘Burst’, with the simulations of the trajectories, denoted by ‘Simul’ on the key. The Fano factor here is 1.99.

Fig. (3) shows histograms obtained from the simulations at the limit of the negative binomial distribution when the bursting takes place. Fig. (3.A) shows a histogram for the time spent by the promoter at the ON state. Fig. (3.B) provides a histogram for the number of mRNA molecules synthesized during each time interval when the promoter is ON. The histogram of the number of mRNA’s after the system has achieved steady state is shown in Fig. (3.C), which gives a comparison of the probability distributions produced by the exact solution (3), a simulation of it by the Gillespie algorithm (10^7^ repetitions) [16], and in the bursting limit (5).

*IV. Discussion.* We provide a biological interpretation of the bursting limit presented in Eq. (5) in terms of the parameter relations that were used. The choice of *A* ∼ 0 implies that the ON→OFF rate *h* ≫ *f*, the OFF→ON rate. Together with the limit *ε, N* ≫ 1, these relationships mean that *k* and *h* are the dominant reaction rates in the bursting limit, and, furthermore, that *h* ≫ *ρ*. Biologically, this means that the promoter ON time is shorter than the mean lifetime of the mRNA’s. In other words, because *h* ≫ *ρ*, most of the mRNA degradation occurs while the promoter is OFF. Also, because *h* ≫ *f*, the promoter tends to be ON for short periods separated by long intervals of OFF. For the short time interval when the promoter is ON, because of the high value of *k*, a large number of mRNA molecules are synthesized. Because mRNA synthesis and ON-OFF switching of the promoter are both stochastic processes, the number of RNA molecules produced in a burst and its time duration are each random variables, and will have different values at each bursting event.

Because *A* is a measure of the proportion of time that the gene is ON, setting *A* very small in the bursting limit is a mathematical expression of the commonsense point that to observe individual bursts, they should be well separated from one another in time. This is apparent in Fig. (1A), where it is evident that 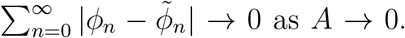 The trajectories of Fig. (2.A) further illustrate this fact. The number of mRNA molecules (Fig. 2.A) increase rapidly during the very short time interval when the promoter is in the ON state, and then decay exponentially after the promoter switches to the OFF state. Figs. (3A) and (3B) are histograms for, respectively, the burst duration and size. Both are random variables arising from intrinsic fluctuations within the cell.

Fig. (1B) shows unimodal probability distributions for high values of *ε* and a range of values of *A*. The distribution of the number of mRNA molecules in this figure are well approximated by a Poisson distribution when *A* is closer 1. In Fig (1B), as *A* → 1, the mode of the distribution moves to the right and its profile approaches that of a Gaussian distribution. The corresponding trajectories of the promoter state and of mRNA numbers are shown on Fig. (2B). The number of mRNA molecules fluctuates around its stationary mean value. The ON and OFF promoter states tend to have durations proportional to *A* along a sufficiently long time interval. The bursts still occur when the promoter is ON, but, because of the slow mRNA synthesis rate in comparison with the duration of the ON states, the trajectories of mRNA numbers in Fig. (2B) have less abrupt increases than Fig. (2A).

Data very similar in appearance to Fig. (2.B) appears in Fig. 4A of a study of transcription driven by the *even-skipped* stripe 2 enhancer in single nuclei of live embryos of *D. melanogaster* [9]. The peaks of differing height were interpreted as individual bursts revealing multiple ON promoter states of differing synthetic capacity through the use of a deterministic model of PolII binding and elongation. The peaks of multiple heights and duration seen in Fig. (2B) shows that the time courses of transcript number observed by Bothma and collaborators [9] can be obtained from two promoter states only.

The two state stochastic model presented here suggests the necessary experimental design for probing the underlying state structure of promoters in *Drosophila* and other organisms. Consider the binary model presented here at the limit of bimodal distributions of *n* as shown on Fig. (1C). The average time intervals for the promoter to be in the ON (*T*_ON_) or OFF (*T*_OFF_) states are similar to each other but longer than the average lifetime of message, *T*_D_ ∼ 1*/ρ* (*T*_ON_∼ *T*_OFF_ *>> T*_D_). Then on average most of the mRNA synthesized during the ON state will be degraded before the promoter switches to the OFF state. When the promoter is OFF, the remaining mRNA will be rapidly degraded. Fig. (2.C) illustrates this fact. In that regime a histogram of the amount of mRNA would be two peaked. The first peak would be near zero and the second peak would be determined by the transcription rate of the ON state. The same reasoning implies that in this regime, an *M* state promoter would produce an *M*-modal histogram.

The experimental realization of this regime depends on the system used. In Ref. [9], observations are made of the fluorescence level of spots of nascent, elongating transcripts. For these experiments, our parameter *ρ* represents the release of completed transcripts from the DNA rather than their physical destruction. Thus, *ρ* would be increased by a shorter probe, albeit at the price of fainter signal. Alternatively or in tandem, an enhancer could be constructed with slower switching between the ON and OFF states in comparison with the rate of transcript elongation and release from DNA. In such conditions multiple underlying ON states would be reflected by multi-peaked histograms, with caveat that any states with extremely fast switching times would be missed.

In the event that the clean experimental regime described above is unobtainable, it is possible that a two state ON-OFF gene could be distinguished from one with multiple states by carefully comparing the durations of productive and degradative periods. Depending on the resolution of the observations, reasonable statistics on the number of mRNA’s synthesized per production event may also be obtained. In both the two state and multistate scenarios, the OFF state (where only degradation occurs) should be exponentially distributed. The distribution of the synthesis period will be dependent on the structure of the underlying promoter states. For example, with a state structure of the form OFF, ON_1_, ON_2_,…, ON_*M*_ where the synthesis rate of ON_*i*_ *<* ON_*i*+1_ and only state transitions that increased or lowered *i* by 1 were permitted, the synthesis times would follow a Gamma distribution. On the other hand, if the observations show a geometric burst size distribution, this favors the bursting limit of the two-state model, given by Eq. (5). However, if the bursting limit does not apply, more complex production distribution may occur [17].

RY was supported by the Natural Sciences and Research Council of Canada and the mobility fellowship CMIRA EXPLORA DOC 2010 of the région Rhône-Alpes (France). AFR and RY would like to thank CAMBAM (at McGill University) for financial support. AFR, GNP, LGS are thankful for CAPES for financial support. JR was supported by NIH Grant No. RO1 OD010936 (formerly RR07801). RY thanks M. Santillán Zerón for useful comments.

